# Yeast display platform for expression of linear peptide epitopes to assess peptide-MHC-II binding in high-throughput

**DOI:** 10.1101/2022.08.09.502759

**Authors:** Brooke D. Huisman, Pallavi A. Balivada, Michael E. Birnbaum

## Abstract

Yeast display can serve as a powerful tool to assess peptide-MHC (pMHC) and pMHC-TCR binding. However, this approach is often limited by the need to optimize MHC proteins for yeast surface expression, which can be laborious and may not yield productive results. Here we present a second-generation yeast display platform for class II MHC molecules (MHC-II) which decouples MHC-II expression from yeast-expressed peptides, referred to as “peptide display”. Peptide display obviates the need for yeast-specific MHC optimizations and increases the scale of MHC-II alleles available for use in yeast display screens. Because MHC identity is separated from the peptide library, a further benefit of this platform is the ability to assess a single library of peptides against any MHC-II. We demonstrate the utility of the peptide display platform across MHC-II proteins, screening HLA-DR, HLA-DP, and HLA-DQ alleles. We further explore parameters of selections, including reagent dependencies, MHC avidity, and use of competitor peptides. This approach presents an advance in throughput and accessibility of screening peptide-MHC-II binding.

## Introduction

Yeast displayed peptide-MHC constructs, originally developed for assessing TCR binding (1–3), have also proven useful for generating large, high-quality datasets on peptide-MHC binding (4–7). In previously described peptide-class II MHC (pMHC-II) yeast display approaches, the peptide is linked to the N-terminus of MHC β chain, which is then expressed in either a βl□1 ‘mini’ (1, 8) or β1β2/□1□2 ‘full length’ format (3, 4, 6, 7). MHC-II proteins have also been expressed on yeast without linked peptide or covalent linkage of MHC □ and β, utilizing leucine zipper fusions to promote heterodimer formation and assessing binding of exogenously added, chemically synthesized peptides (5). An additional approach expresses a secreted MHC-II in yeast for co-display with a yeast surface-expressed peptide (9). While theoretically any MHC allele can be analyzed by yeast display, the relatively simple protein folding machinery in yeast can cause some MHCs, especially MHC-IIs, to not fold properly despite detectable surface expression (1, 4, 8). This necessitates each MHC be validated for fold, such as through the binding of a recombinantly expressed TCR (1, 8). In the absence of detectable TCR binding, error-prone mutagenesis of the MHC is required to identify mutations that can stabilize the MHC fold while not affecting the peptide- and receptor-binding surfaces (1, 8). This process can be time and effort intensive, has uncertain success, and requires availability of reagents such as a known MHC-restricted TCR.

To circumvent the need for individual optimization of MHC proteins, we have developed a second-generation yeast display approach for assessing pMHC-II binding which does not require yeast-specific MHC-II optimizations. In this approach, peptide and MHC-II expression are decoupled, with a library of peptides displayed on the surface of yeast and MHC-II protein expressed as recombinant soluble molecules. Because of this decoupling and exclusive presentation of peptides on the yeast surface, we refer to this approach as “peptide display”.

Here, we outline the system, explore experimental conditions, and compare the results with existing yeast display datasets. We apply this approach to study HLA-DR alleles, followed by HLA-DP and HLA-DQ alleles, which have to this point been largely inaccessible in previous yeast display approaches due to few HLA-DP or HLA-DQ alleles being validated as folded on the yeast surface (5). Existing pMHC yeast display platforms further require generation of a new library for each new MHC being screened (1, 2, 4, 6–8). In contrast, because the peptide library is decoupled from MHC, a single peptide display library is extensible to any MHC-II without requiring additional library cloning and generation, expanding the utility of the approach.

## Results

### Design of peptide display libraries and experiments

For the peptide display approach, we adapted the pCT302 yeast display vector, which fuses peptide sequences of interest to the C-terminus of the yeast mating protein Aga2 (10). We left the N-terminus of the construct unchanged, which contains Aga2 fused to an epitope tag and a subsequent C-terminal linker sequence. To the C-terminus of this linker, we connected the N-terminus of a peptide sequence of interest. In order to readily assess frameshift mutations in our peptide sequence, we linked the displayed peptide’s C terminus to a Myc epitope tag via a short linker.

To examine MHC-peptide repertoires, MHC-II proteins were expressed recombinantly rather than co-expressing MHC-II proteins on the surface of yeast, which circumvents yeast protein folding concerns. MHC-II proteins were stabilized with a linked portion of class II-associated Ii peptide (CLIP_87-101_, PVSKMRMATPLLMQA) (11), connected to the N-terminus of MHC-IIβ chain via a linker containing a 3C protease cleavage site. The MHC-II protein was site-specifically biotinylated, allowing for easy functionalization and visualization with fluorescent streptavidin (SAV).

To assess pMHC-II binding, yeast were co-incubated with fluorescent tetramerized MHC-II protein, 3C protease, and HLA-DM (**Figure 1A**). 3C protease cleaves the CLIP-MHC linker, enabling the stabilizing peptide to dissociate. The peptide exchange catalyst HLA-DM is available to assist with dissociation of CLIP and binding to the yeast-expressed peptide (12). If the MHC is able to bind to the peptide of interest, we can detect fluorescence via flow cytometry-based assessment of the yeast population, akin to a primary antibody stain. When we stain peptide-displaying yeast with soluble tetramerized HLA-DR401, we observe a clear fluorescent population for yeast expressing a version of an Influenza A HA_306-318_ peptide (APKYVKQNTLKLAT) known to bind HLA-DR401(HLA-DRA1 *01:01 / DRB1*04:01). (Hennecke and Wiley, 2002; Rappazzo et al., 2020), with minimal binding to an off-target CD48_36-51_ peptide (FDQKIVEWDSRKSKYF) (**Figure 1B**). CD48_36-51_ peptide binds to a related allele, HLA-DR402 (HLA-DRA1*01:01 / DRB1*04:02) (4), but is likely dis-preferred for HLA-DR401 binding because of the large aromatic Trp residue at the expected P4 pocket in the optimal register.

**Figure 1.**
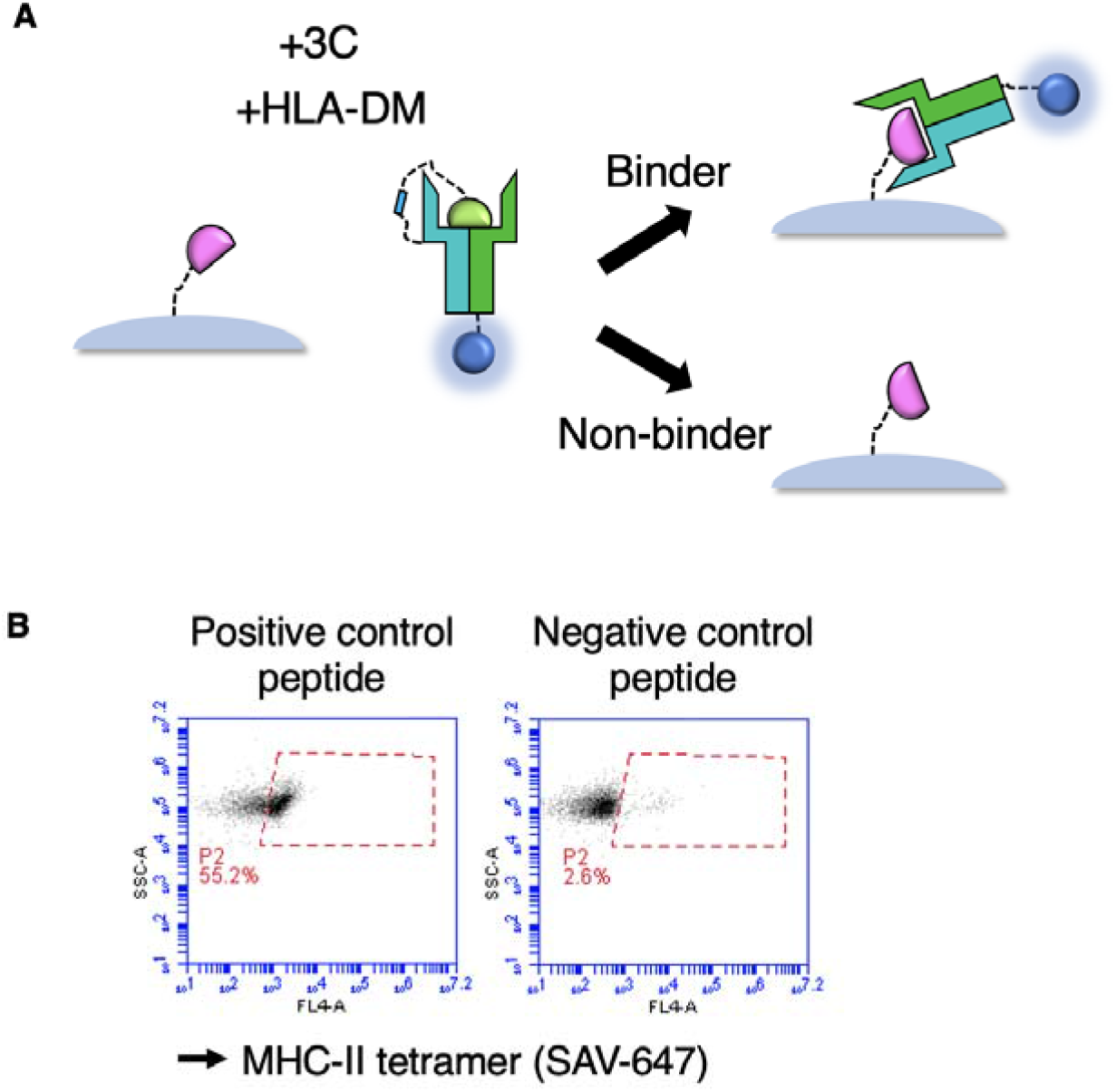
Overview of peptide display formatting and experiments. **A)** Peptide display platform formatting and selection strategy. Peptides are expressed on the surface of yeast, and MHC is expressed recombinantly, with CLIP_81-101_ peptide (green) for stability. Upon addition of HLA-DM and 3C protease to cleave the CLIP-MHC linkage, the MHC can bind to yeast-expressed peptides (pink). Binders can be manipulated using a handle on the MHC, such as a fluorescent streptavidin. **B)** Flow cytometry analysis of tetramerized HLA-DR401 with binder (HA-derived) and non-binder (CD48-derived) peptides.

To explore the constraints of the system and reliance on each component of the selection reaction mix, we separately titrated HLA-DR401 tetramers, HLA-DM, and 3C protease and assessed pMHC-II binding. As expected, there was an MHC tetramer concentration dependence, with binding signal decreasing as MHC concentration decreased (**Supplemental Figure 1A**). We similarly observe a dependence on the presence of HLA-DM, with largely unchanged signal across concentrations from 0.5 μM to 3.5 μM, and with signal lost at the lowest concentration tested, 75 nM (**Supplemental Figure 1B**). Next, we assessed if the peptide exchange reaction could proceed without cleaving the linker between recombinant MHC-II and the stabilizing CLIP peptide. We observe signal is lost when we decrease 3C concentrations, illustrating a need to allow the stabilizing peptide to freely dissociate in order to enable loading of the yeast-expressed peptide (**Supplemental Figure 1C**; **Figure 1C** is a subset of this figure). Across all experiments, the negative control peptide showed minimal signal at reagent concentrations tested. From these experiments, optimized reagent concentrations were selected for utilization in large-scale experiments (75 nM MHC-II tetramer, 0.5μM HLA-DM, and 1 μM 3C).

### Application of peptide display platform to screen HLA-DR401

Next, to utilize the peptide display approach for screening peptides at a repertoire-scale, we generated a library of randomized 13mer peptides, containing approximately 5 x 10^6^ unique peptides. We performed selections with HLA-DR401, which would enable comparison with existing pMHC-II datasets (4, 13). Yeast were incubated with fluorescent tetramerized HLA-DR401, 3C protease, and HLA-DM in phosphate buffered saline (PBS) for 45 minutes. The neutral pH allows 3C protease to cleave the linker which connects MHC-II to the stabilizing CLIP peptide. To mimic acidic endosomal conditions, yeast were then incubated with the reaction mixture in pH 5 saline for an additional 45 minutes. Next, yeast were incubated with anti-fluorophore magnetic beads. Yeast which expressed peptides that bound to HLA-DR401 were enriched utilizing a magnetic enrichment strategy.

We performed three iterative rounds of selection, growing yeast to confluence between each round. In each round, we sampled yeast after the first incubation in PBS, the second incubation in acidic saline, and after elution from the magnetic column and examined the fluorescent population via flow cytometry. Over subsequent rounds, the population of fluorescent MHC-bound yeast increased at each of these steps (**Supplemental Figure 2**).

Next, we determined the identities of the peptides enriched for binding to HLA-DR401 by deep sequencing each round of the library selections. Given the open MHC-II binding groove and 13mer length of peptides assessed, we inferred the binding register of each peptide utilizing a position weight matrix approach (7). Peptides inferred to bind using the central register are represented in **Figure 2**. These representations include fractions of peptides with amino acids at each position (**Figure 2A**), log_2_ fold change compared to the unselected library (**Figure 2B**), and sequence logos (**Figure 2C**). We observe a strong P1 preference for large hydrophobic and aromatic hydrophobic residues; a preference at P4 for acidic residues and non-aromatic hydrophobic residues; a preference for polar residues at P6; and preference for small residues at P9. The auxiliary anchor P7 exhibits some preference for Pro, Leu, and Asn. This motif is consistent with motifs from other high-throughput datasets (4, 13–16), with a greater representation of P4 hydrophobic residues relative to acidic residues, though peptides with hydrophobic P4 residues have been shown to bind to HLA-DR401 (4). Using the peptide display approach, we were able to enrich for peptides which bind to HLA-DR401, which are consistent with previously described motifs and examples of HLA-DR401-binding peptides.

**Fig 2.**
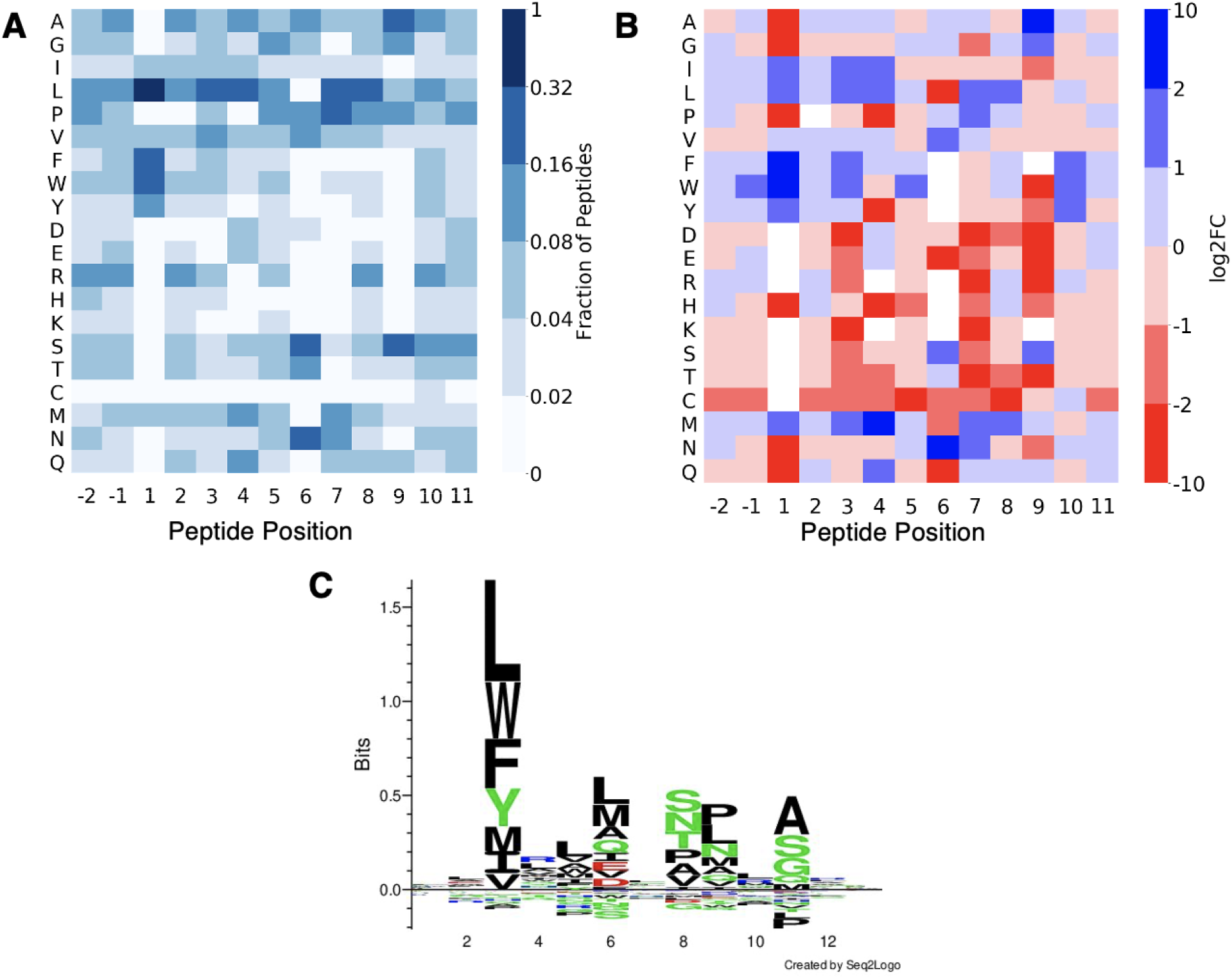
Motifs of enriched peptides in the central register from HLA-DR401 tetramerbased library selections. **A)** Heatmap highlighting the fraction of peptides with each amino acid at each position. **B)** Log_2_ fold change of positional amino acid frequency compared to unselected library. Amino acids which did not appear in the enriched sequences at a given position are white. **C)** Sequence logo of enriched peptides.

We next assessed binding to a panel of individual peptides with known affinity values for HLA-DR401 (4). Performing binding reactions similar to selection experiments, we utilized flow cytometry to examine binding after the first incubation in PBS and the second incubation in acidic saline (**Figure 3**). After the incubation in acidic saline, we observe a clear monotonic relationship between measured IC_50_ value and percent MHC-tetramer positive population or mean fluorescence intensity. This relationship is only clear after the second incubation, which suggests that the acidic incubation may be helpful in enabling pMHC-II binding or establishing the protonation state of acidic residues. The clear binding of peptides containing acidic residues also suggests that acidic residue-containing peptides, including peptides with acidic P4 residues, are not systemically excluded from MHC binding in this format at an individual peptide level.

**Figure 3.**
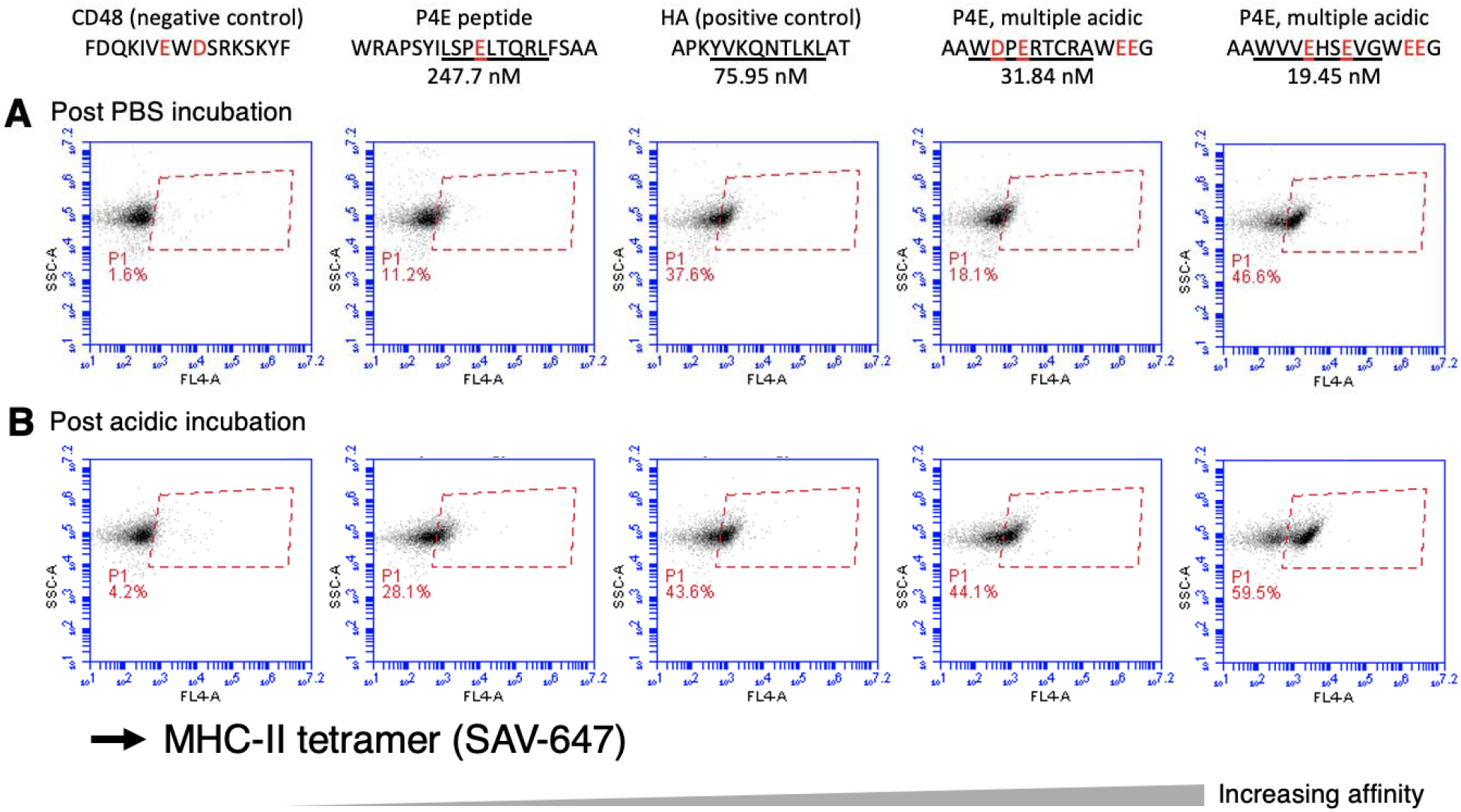
Tests on clonal yeast populations containing acidic residues. HLA-DR401 tetramer staining on individual yeast clones after **A)** incubation in pH 7.2 PBS and **B)** subsequent incubation in acidic saline. CD48-derived peptide was included as a negative control and remaining peptides have previously measured affinities indicated. HA peptide affinity was previously measured with an additional C-terminal Gly, which is present in the peptide display C-terminal linker. Central 9mer cores from inferred registers, based on similarity with HLA-DR4O1 peptide motifs, are underlined for each peptide, and acidic residues highlighted in red.

To explore the requirement for MHC avidity and to examine alternative modes of selections, we performed selections with monomeric MHC. Performing monomeric selections also provides us a mode to assess if charged AlexaFluor647, which is conjugated to our streptavidin, could be affecting P4 acidic preference. Motifs from monomer selections (**Supplemental Figure 3**) are largely similar to the tetrameric selections (**Figure 2**), suggesting MHC avidity in tetrameric selection is not necessary and the use of streptavidin-AlexaFluor647 (SAV-647) is not biasing peptide binding.

### Assessing additional MHCs, including HLA-DP and HLA-DQ alleles

Given the clear enrichment with HLA-DR401 we set out to demonstrate the extensibility of this approach to other MHC proteins by performing selections for HLA-DQ0602 (HLA-DQA1*01:02 / DQB1*06:02), HLA-DQ0603 (HLA-DQA*01:02 / DQB1*06:03), HLA-DP401 (HLA-DPA1*01:03 / DPB1*04:01), and HLA-DR15 (HLA-DRA1*01:01 / HLA-DRB1*15:01). These proteins represent alleles expressed by each of the three canonical MHC-II genes (17). Further, these alleles are implicated in autoimmunity, with HLA-DQ0602 and HLA-DR15 comprising a haplotype linked to protection from type 1 diabetes, as well as susceptibility to multiple sclerosis (18–20).

For all four alleles, we see clear enrichment, with preferences at MHC anchor residues (**Figure 4**). HLA-DR15 prefers hydrophobic residues at P1; aromatic residues at P4; Asn, Ser, and Gly at P6; and small hydrophobic Leu, Val, Ala at P9 (**Figure 4A**). These preferences closely resemble previously reported motifs (13, 14). HLA-DP401 has strong preferences at P1, P6, and P9 for hydrophobic residues, including aromatics at P1 and P6. This motif is similar to those in existing datasets (13, 14). However, while somewhat present and most visible in the heatmaps (**Figure 4B**), the preference for P4 Glu, Thr, and Ser is less pronounced than in the mass spectrometry comparison datasets (13, 14). HLA-DQ0602 has clear preference at P1, P4, P6, P7, and P9, alongside greater amino acid preference at non-anchors compared to the other alleles studied here (**Figure 4C**). The anchor preferences and increased preference across the peptide is generally consistent with previously reported peptides from mass spectrometry (13). The peptide display approach can also elucidate differences between closely related alleles. A related HLA-DQ allele, HLA-DQ0603, showed similar P1, P4, and P9 preferences to HLA-DQ0602. The beta chain of HLA-DQ0603 differs from the HLA-DQ0602 beta chain at two amino acids, Phe9Tyr and Tyr30His, which affect P6 and P7 preferences (**Supplemental Figure 4A**, **Figure 4C-D**) (21). HLA-DQ0603 exhibits a greater preference for P6 Asp and Ser, likely due to the positive His at HLA-DQ beta position 30 (**Supplemental Figure 4B**, adapted from PDB 6DIG (22)). Similarly, these polymorphisms generate a distinctive preference for Pro at the auxiliary P7 anchor, which is not captured in the NetMHCIIpan4.0 motif viewer (16). These data demonstrate the utility of the peptide display approach for extending yeast display assessment of peptide-MHC binding across alleles, especially alleles not previously validated on yeast.

**Figure 4.**
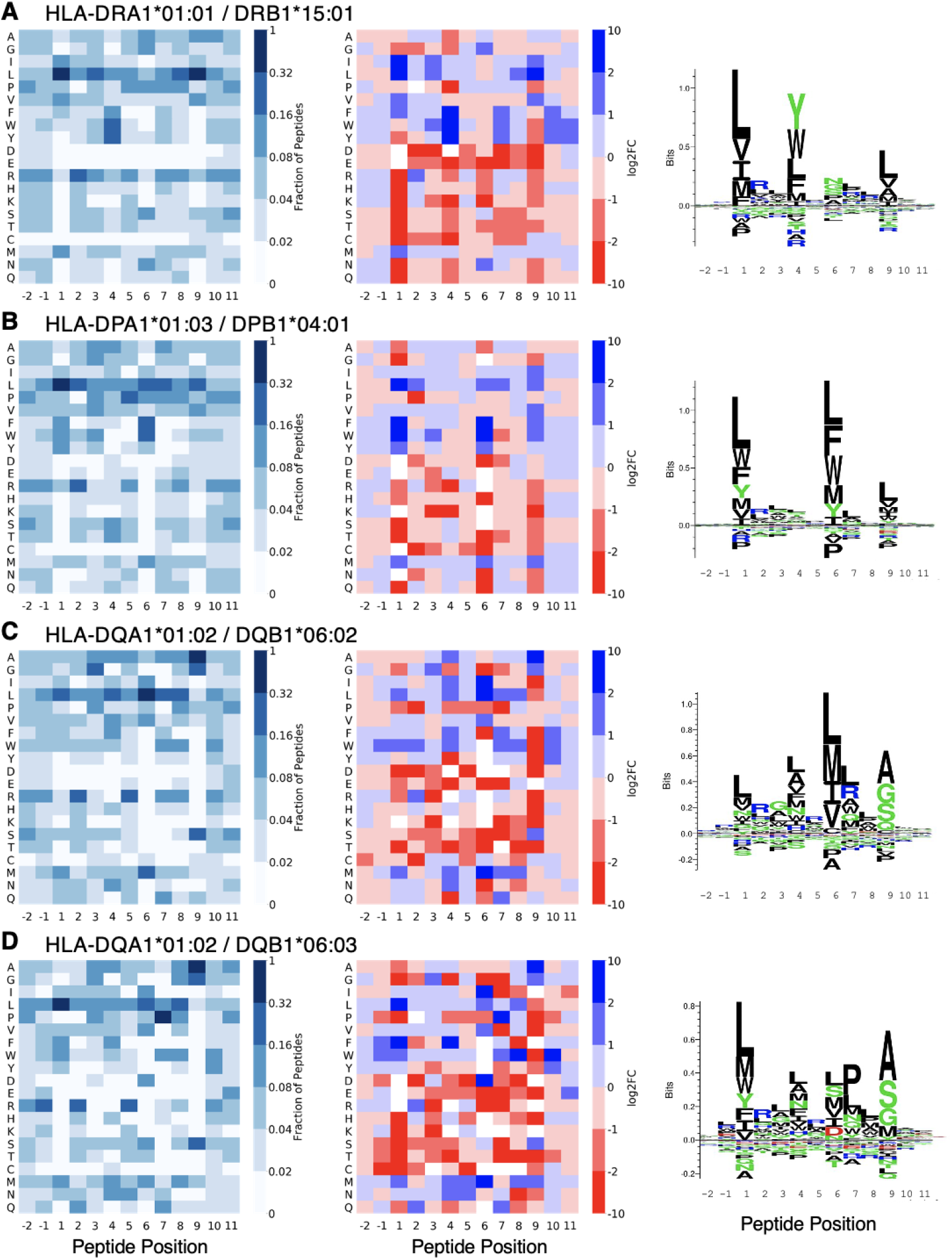
Motifs for peptides enriched for additional MHC-II alleles. Motifs for **A)** HLA-DRB1*15:01 (HLA-DR15), **B)** HLA-DPA1*01:03 / DPB1*04:O1 (HLA-DP401), **C)**HLA-DQA1*01:02 / DQB1*06:02 (HLA-DQ0602), and **D**) HLA-DQA1*01:02 / DQB1*06:03 (HLA**-**DQ0603). **Left:** Heatmap highlighting the fraction of peptides with each amino acid at each position. **Center:** Log_2_ fold change of positional amino acid frequency compared to unselected library. Amino acids which did not appear in the enriched sequences at a given position are white. **Right:** Sequence logo of enriched peptides.

### Adding exogenous peptide as a competitor

The peptide display approach is highly tunable across multiple parameters. As an example, one such tunable parameter is the addition of competitor peptide of known affinity. To examine the effects of adding exogenous peptide on MHC binding to yeast-expressed peptides, we titrated competitor binding and assessed its effects on peptide-MHC-II binding (**Supplemental Figure 5**). Our exogenous peptide was used to separately compete with two peptides expressed on the surface of yeast. We expressed CLIP_81-101_ peptide (LPKPPKPVSKMRMATPLLMQA) on yeast, as well as a synthetic HLA-DP401 binder which was the most frequent peptide in the inferred central binding register from our HLA-DP401 selections (KFFLLLPMCVWCK). The synthetic strong binder matches with the overall motif of enriched peptides, with hydrophobic anchor residues. We assessed binding of HLA-DP401 to each of the yeast-expressed peptides in the presence of varying concentrations of exogenously added CLIP_81-101_ peptide. We see a clear loss of MHC tetramer signal as we increase the concentration of competitor peptide, with the synthetic strong binder retaining signal at higher concentrations of competitor than the yeast-expressed CLIP peptide. Addition of exogenous peptide, such as a weak and pan-MHC-II allele binder like CLIP (23), presents an opportunity to further segregate weak and strong binders in selections and another method for tuning stringency of selection experiments.

## Discussion

Yeast-displayed peptide-MHC constructs have proven useful for assessing peptide-MHC and pMHC-TCR binding, but have been is limited by the need to individually validate and optimize MHCs for yeast expression (1, 4, 8). To overcome this challenge, we have developed a yeast surface display platform that decouples peptide and MHC expression, enabling users to express MHCs without optimizing them for yeast expression. Libraries of peptides are expressed on the surface of yeast, and binding to recombinantly-expressed MHCs is assessed. Previous yeast display approaches decoupling peptide and MHC expression have expressed both peptide and MHC in yeast (9), which still presents the limitations of yeast protein folding, whereas expressing MHC-II proteins in an orthogonal expression system such as insect cells enables more reliable MHC-II protein folding. Separating the MHC from yeast expression also enables users to make a single library which can be screened against any recombinantly expressed MHC-II protein, so assessing a new MHC does not require generation of a new yeast display library. Here we demonstrated the principle of the platform (**Figure 1**) and have shown utility for generating datasets across HLA-DR, HLA-DP, and HLA-DQ alleles (**Figure 2**, **Figure 4**). Using a single peptide library and expressing these MHCs recombinantly, we were able to quickly generate high quality, large-scale datasets of peptide binders.

Peptide-MHC-II binding in the peptide display platform captures a composite of binding dynamics. Namely, peptide display requires dissociation of the invariant CLIP peptide from the MHC groove and binding of other peptides. As a result, we can capture the interplay of CLIP dissociation and peptide binding to MHC, and potential equilibrium binding and unbinding. In contrast, many previous yeast display library profiling systems capture only peptide dissociation since peptides of interest begin bound to the MHC-II (3, 4, 7). Additionally, the clear dependence on HLA-DM presence (**Supplemental Figure 1B**) stands in contrast to the minimal HLA-DM reliance in a dissociation-based pMHC-II platform, which has previously shown similar motifs in library selections with or without HLA-DM (4), suggesting HLA-DM may be required for dissociation of CLIP and association of test peptides.

Selection strategies using peptide display are versatile and tunable. For instance, we demonstrate the use of tetramerized MHC-II proteins (**Figure 2**) as well as monomeric protein for selections (**Supplemental Figure 3**), and the use of exogenous competitor peptide for tuning binding stringency (**Supplemental Figure 5**). Further, there is great flexibility for use of MHC molecules. Essentially any source of recombinantly expressed class II MHCs with exchangeable peptides can be utilized, including commercial sources used for tetramers (24). Production should be agnostic to use of bacterial (25), insect (1), or other mammalian expression systems (26, 27). We also explored constraints of the system, including reliance on 3C protease, HLA-DM, and acidic conditions (**Supplemental Figure 1** and **Figure 3**). Further exploration of the construct itself can help tune the platform for downstream applications and ensure minimal construct-specific artifacts.

The peptide display platform may impact multiple application areas. Data from high-throughput yeast display screens have shown benefit for training prediction algorithms (4), and the generalizability of the peptide display platform across MHC alleles can help expand data generation potential. Additionally, given the extensibility across alleles and possibility to combine with defined peptide libraries (6, 7), this platform would be ideal for generating and assessing binding of whole human or bacterial proteomes to MHC and, potentially pMHC complexes to TCR, for applications in antigen discovery in infectious disease, autoimmunity, cancer, and transplantation.

### Experimental Procedures

#### Vector formatting for peptide display

pCT302 vector (10) was adapted to fuse a peptide sequence to Aga2. We left unchanged the N-terminus of the construct, in which Aga2 is connected via a linker to an HA epitope tag (YPYDVPDYA) (10). We also leave unchanged the subsequent C-terminal linker sequence (LQASGGGGSGGGGSGGGGSAS) (10). The C-terminus of this linker is connected to the N-terminus of our peptide sequence of interest. To assess frameshift mutations in our peptide sequence, we link the peptide to a Myc epitope tag (EQKLISEEDL) via a short linker (GGSGG). The Myc tag is proceeded by a stop codon. pCT302 was a gift from Dane Wittrup.

#### Peptide display parameter exploration experiments

To optimize peptide display reaction conditions, we generated reaction mixtures containing MHC-II tetramers, HLA-DM, and 3C protease in PBS. Yeast were washed with PBS then resuspended in the reaction mixture made in PBS. Yeast were incubated for 45 minutes at room temperature, rotating, then washed into FACS buffer (0.5% BSA and 2 mM EDTA in 1x PBS) and assessed via flow cytometry (Accuri C6 flow cytometer, BD Biosciences; Franklin Lakes, New Jersey). In the reaction mixture for MHC-II titration, we included 1 μM 3C protease and 3.5 μM of HLA-DM. In the reaction mixture for HLA-DM titration, we included 1 μM 3C protease and 75 nM MHC tetramer. And in the reaction mixture for 3C protease titration, we included 0.5 μM HLA-DM and 75 nM MHC tetramer.

For testing individual peptides with measured affinity values (**Figure 3**), similar conditions were utilized: 75 nM HLA-DR401 tetramer, 1 μM 3C, and 0.5 μM HLA-DM. Yeast were also subsequently incubated in a nine-fold excess of acidic saline and sampled after 45 minutes. Experiments assessing the impacts of CLIP as a competitor peptide (**Supplemental Figure 5**), utilized the same concentrations, with 75 nM HLA-DP401 tetramer, 1 μM 3C, 0.5 μM HLA-DM, in addition to 10 μM TCEP and 20-minute pre-incubation of yeast with TCEP-containing PBS before incubating with the reaction mixture. Yeast were sampled after a 45-minute room temperature incubation.

MHC tetramers were generated using streptavidin-AlexaFluor647 (SAV-647; made in-house as previously described (28)), with a 5-fold molar ratio of MHC to SAV-647, which has four binding sites. Tetramer concentration is given as the concentration of SAV-647, representing the concentration of tetrameric species.

#### Randomized library generation

A 13mer randomized peptide library was generated by performing polymerase chain reaction (PCR) with a degenerate primer encoding NNK codons (N = A, C, T, or G; K = G or T). We utilized a stop codon-containing template for PCRs to decrease the likelihood of contamination with a strong-binding construct. Libraries were generated by electroporating RJY100 yeast (29) with a 5:1 mass ratio of randomized peptide product and linearized pCT302 vector.

#### Recombinant protein expression

Proteins were expressed recombinantly, as described previously (4, 7). The extracellular domains of the α and β chains from HLA-DR401, HLA-DR15, HLA-DQ0602, HLA-DQ0603, HLA-DP401, and HLA-DM were cloned into pAcGP67a vectors for expression in insect cells. The ectodomains of each chain were formatted with a poly-histidine purification site and encoded on separate plasmids. The α chains contained an AviTag biotinylation site (GLNDIFEAQKIEWHE). Excepting HLA-DM, the N-termini of β chains were linked to CLIP_81-101_ peptide via flexible Gly-Ser linker containing a 3C protease site (LEVLFQGP). Plasmid DNA was transfected into SF9 insect cells with BestBac 2.0 DNA (Expression Systems; Davis, CA) using Cellfectin II reagent (Thermo Fisher; Waltham, MA). Viruses for α and β chains were co-titrated to determine an optimal ratio for heterodimer formation, co-transduced into High Five (Hi5) insect cells (Thermo Fisher), and incubated for 48-72 hours. Proteins were secreted by Hi5s and purified by conditioning the cell culture media with 50 mM Tris-HCl (pH 8), 1 mM NiCl_2_, and 5 mM CaCl_2_, clearing the precipitant via centrifugation, and incubating the resulting supernatant with Ni-NTA resin (1). Proteins were further purified using a S200 increase column on an AKTAPURE FPLC (GE Healthcare; Chicago, IL). Excepting HLA-DM, protein was site-specifically biotinylated and confirmed using a gel-shift assay (30).

#### Library selections

The randomized 13mer library had 1 x 10^8^ unique transformants and a random subsampling of 5 x 10^6^ yeast, based on optical density, were grown up for use in selections in order to increase coverage of enriched yeast by sequencing and for easier comparison across iterations of selections. An overrepresentation of these yeast were input into Round 1, with 5 x 10^7^ yeast as input, measured by optical density. Rounds 2 and 3 used 2.5 x 10^7^ – 5 x 10^7^ input yeast.

To perform tetrameric selections, yeast were first washed into pH 7.2 PBS with 10 μM TCEP and incubated for 20 minutes while making the reaction mixture. The reaction mixture contained MHC tetramers, generated by pre-incubating 75 nM SAV-647 and 375 nM biotinylated MHC. Tetramers are added to the appropriate volume of pH 7.2 PBS with 10 μM TCEP. This is followed by the addition of 3C protease to 1 μM, then the addition of 0.5 μM HLA-DM (excepting Round 3 of HLA-DR401 no-streptavidin selections, which received 1 μM HLA-DM). TCEP was added to prevent disulfide bonds which could diminish the enrichment of peptides containing cysteine. Yeast were spun down and resuspended in reaction mixture at 2 x 10^8^ yeast/mL. This reaction was incubated at room temperature for 45 minutes, rotating, covered to decrease risk of photobleaching. Then, yeast in the reaction mixture were placed on ice, and a subsample taken for assessment via flow cytometry, and the remaining yeast were moved to a nine-fold excess of pH 5 citric acid saline buffer (20 mM citric acid, 150 mM NaCl). Yeast were incubated in the acid saline at 4°C for 45 minutes, rotating. Yeast were then washed into FACS buffer, again at 2 x 10^8^ yeast/mL, and ∝-AlexaFluor647 beads (Miltenyi Biotec, Bergisch Gladbach, Germany) equal to 10% of the FACS buffer volume were added. Yeast were incubated with beads at 4°C, rotating for 15 minutes. Yeast were then pelleted, resuspended in 5 mL FACS buffer, and run over a Miltenyi LS column (Miltenyi Biotec), washed three times with 3 mL FACS buffer, and eluted with 5 mL FACS buffer. The elution contained the yeast which bound to the MHC. Selected yeast were then washed into SDCAA media and grown to confluence. Yeast were sub-cultured in SGCAA media at OD_600_ = 1 for 48-72 hours at 20°C before each experiment (31).

Monomeric selections were performed similarly to tetrameric selections, with a few modifications. Monomeric MHC-II was added in the reaction mixture, absent SAV-647. 300 nM MHC-II was added, equivalent to the amount of MHC-II which binds to the four binding sites of 75 nM SAV-647 in tetrameric selections. Streptavidin beads (Miltenyi Biotec) were added in place of ∝-AlexaFluor647 beads, and selections proceeded in the same manner.

#### Deep sequencing and data processing

Peptide identities were determined from enriched yeast following selections. Plasmid DNA was extracted from 5 x 10^7^ input yeast via the Zymoprep Yeast Miniprep Kit (Zymo Research; Irvine, CA) following manufacturer’s instructions. Amplicons were generated by two rounds of PCR to add inline i5 and i7 paired-end handles and inline sequencing barcodes, unique to rounds multiplexed in a single sample. PCR primers were designed to capture the 13mer peptide sequence, priming off of adjacent constant sequences. Amplicons were sequenced with an Illumina MiSeq (Illumina Incorporated; San Diego, CA) at the MIT BioMicroCenter using 300 nucleotide kits.

Paired-end reads were assembled using PandaSeq (32). Given the large space of possible NNK-encoded 13mer peptides, single nucleotide variants are likely to be a result of PCR or sequencing errors. Using CDHIT (33), single nucleotide-variants were combined and the more frequent sequence chosen as the consensus sequence.

#### Peptide binding register inference and motif visualization

To infer the register in which the enriched 13mer peptides bound, we utilized a position weight matrix (PWM) method (7). Enriched peptides from round 3 of selection were one-hot encoded and padded on the C-terminus of the peptide. We utilized a 9mer PWM and eight registers and one ‘trash’ register.

At the start, peptides were randomly assigned to registers and a 9mer PWM was generated. The contribution of a given peptide to the PWM was weighted by its read count. Over successive iterations, peptides were assigned to new registers and the PWM updated, excluding peptides in the trash cluster. Assignment to each register is stochastic, but biased, such that registers which match the PWM were favored. At each assignment, if the peptide was assigned to a non-trash register, we first took out that peptide from the PWM. The PWM then defined an energy value for each register option, which was used to generate a Boltzmann distribution from which the updated register shift was sampled. Stochasticity was decreased over time by raising the inverse temperature linearly from 0.05 to 1 over 20 iterations, simulating cooling (34). The final iteration was deterministic, where the distribution concentrated solely on the locally optimal register option.

From selections with HLA-DR401 (tetramers), HLA-DR401 (monomers), HLA-DP401, HLA-DR15, HLA-DQ0602, HLA-DQ0603 we identify 11,275 / 32,885 / 89,746 / 46,276 / 27,231 / 8,289 enriched peptides in our sequencing after Round 3, respectively. Of these, from register-inference, we identify 6,164 / 18,845 / 56,977 / 28,167 / 11,584 / 2,219 peptides in native registers, respectively.

Heatmap and sequence logo visualizations were generated for peptides which were inferred to bind in the central register, with two peptide flanking residues on either side of the 9mer core. Heatmaps were generated using a custom script to show amino acid frequency or log_2_ fold change over the unselected library at each position. Amino acids which did not appear at a given position in the round 3 library appear as white spaces in the log2 fold change heatmap. Sequence logos were generated using Seq2Logo (35), with default settings, except background frequencies were generated from the unselected library. In heatmaps and sequence logos visualizations, peptides were not weighted by read counts.

## Data Availability

All deep sequencing files are deposited on the Sequence Read Archive (SRA) with accession code PRJNA867335 (http://www.ncbi.nlm.nih.gov/bioproject/867335). Round 3 enriched peptides with stop codon-containing peptides removed are available in **Supporting Information 1**. Inferred register clusters for enriched peptides are indicated, with the trash register indicated as register “-1”. Peptides from the unselected library are also provided in **Supporting Information 1**.

## Code Availability

Scripts used for data processing, register inference, and visualization are publicly available at https://github.com/birnbaumlab/Huisman-et-al-2022-Peptide-Display.

## Acknowledgements

We would like to thank Zheng Dai and David Gifford for helpful discussions and thoughtful review of the manuscript. Thank you to the MIT BioMicro Center for performing Illumina library sequencing. This work was supported in part by the Koch Institute Support (core) Grant P30-CA14051 from the National Cancer Institute. This work was supported by National Institute of Health (U19-AI110495), the Melanoma Research Alliance, the Packard Foundation, and a Schmidt Futures grant to MEB, and a National Science Foundation Graduate Research Fellowship to BDH.

## Author Contributions

Project conception: B.D.H and M.E.B.; Conducted experiments, B.D.H and P.A.B.; Performed Data Analysis: B.D.H. and P.A.B.; Supervision: M.E.B.; Wrote manuscript: B.D.H. and M.E.B.; all authors edited the manuscript.

