## Supplemental Information for "Yeast display platform for expression of linear peptide epitopes to assess peptide-MHC-II binding in high-throughput"

**5 Supplemental Figures**

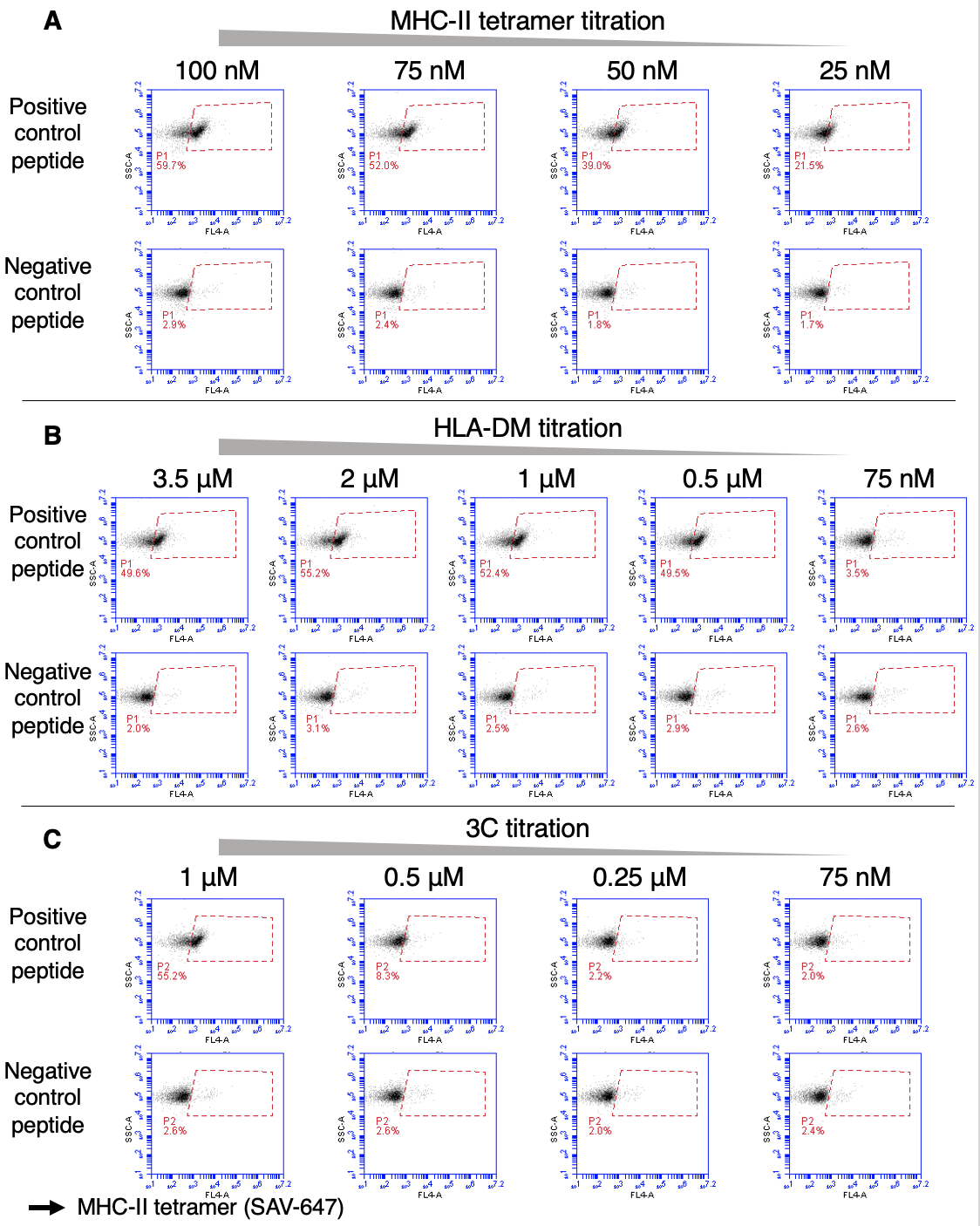

**Supplemental Figure 1. Testing experimental conditions.** Reliance on each component of the selection reaction mix is assessed with titrations of **A)** HLA-DR401 tetramers, **B)** HLA-DM, and **C)** 3C protease with binder (HA-derived) and non-binder (CD48-derived) peptides.

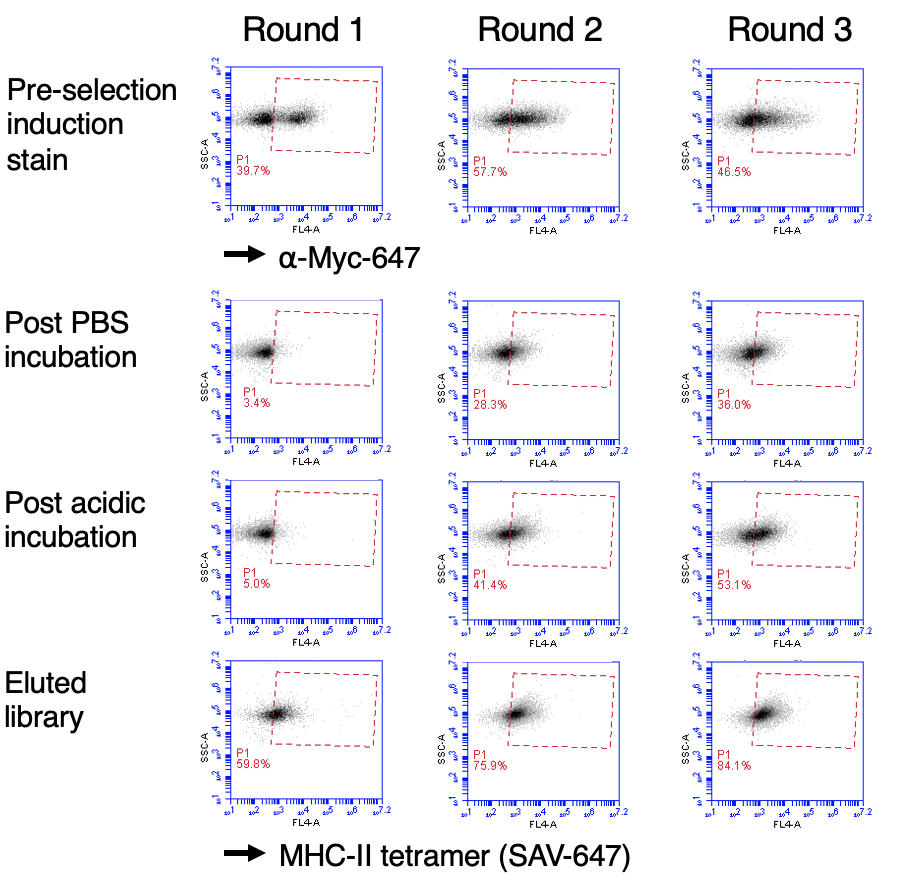

**Supplemental Figure 2. Dynamics of library selections for HLA-DR401 tetramer-based library selections.** Pre-selection induction staining with anti-epitope tag antibody, as well as tetramer staining at timepoints throughout each selection, including after incubation in pH 7.2 PBS, after subsequent incubation in acidic saline, and after elution from magnetic enrichment column.

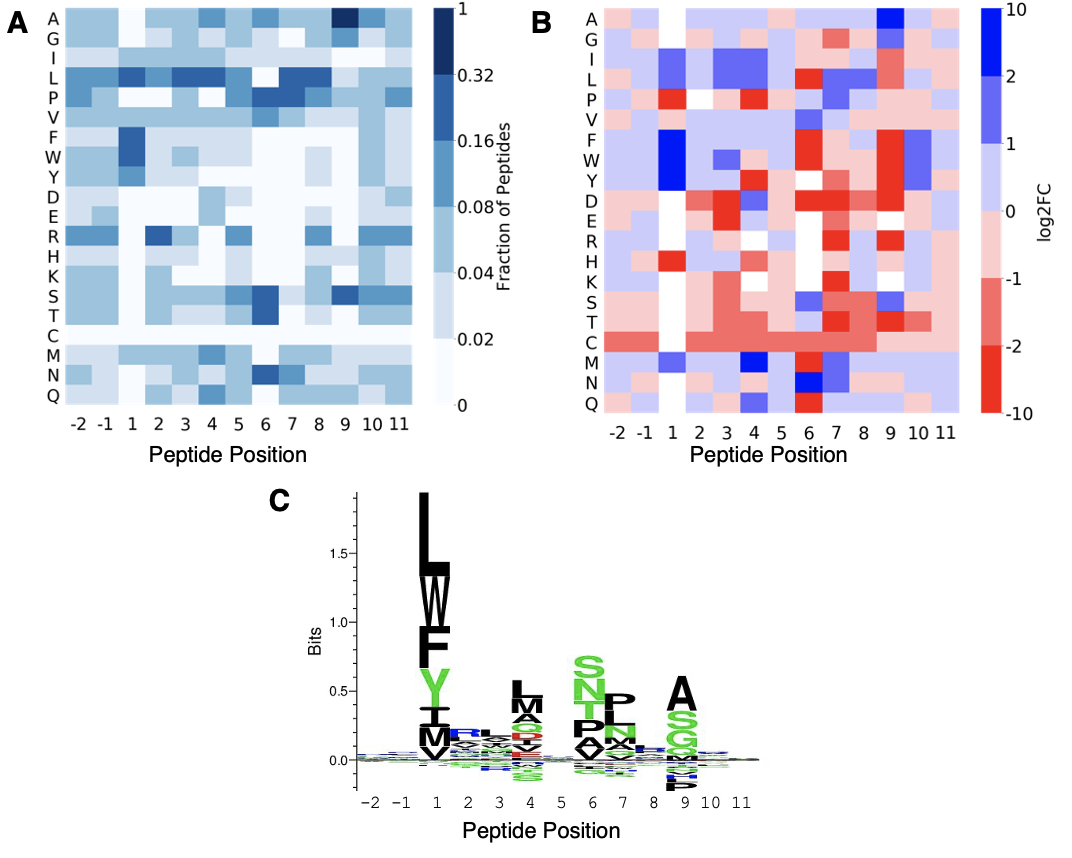

**Supplemental Figure 3. Motifs from library selections with HLA-DR401 monomers without fluorescent streptavidin. A)** Heatmap highlighting the fraction of peptides with each amino acid at each position. **B)** Log_2_ fold change of positional amino acid frequency compared to unselected library. Amino acids which did not appear in the enriched sequences at a given position are white. **C)** Sequence logo of enriched peptides.

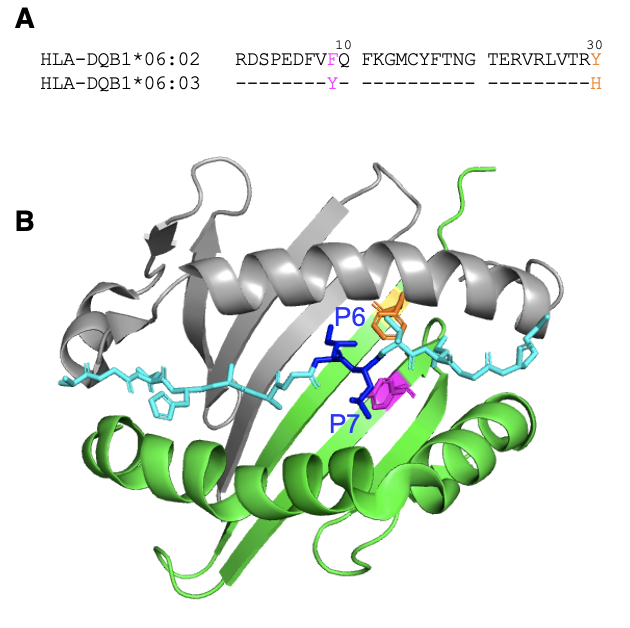

**Supplemental Figure 4. Related HLA-DQ alleles. A)** Sequence alignment of related HLA-DQ alleles assessed in this study: HLA-DQB1*06:02 and HLA-DQB1*06:03, with polymorphisms highlighted. **B)** Structure of HLA-DQA1*01:02 (grey) / HLA-DQB1*06:02 (green) with peptide (cyan), adapted from PDB 6DIG. Residues that differ between HLA-DQB1*06:02 and HLA-DQB1*06:03 are highlighted in magenta and orange, as in **A**, and interacting peptide residues P6 and P7 are in blue.

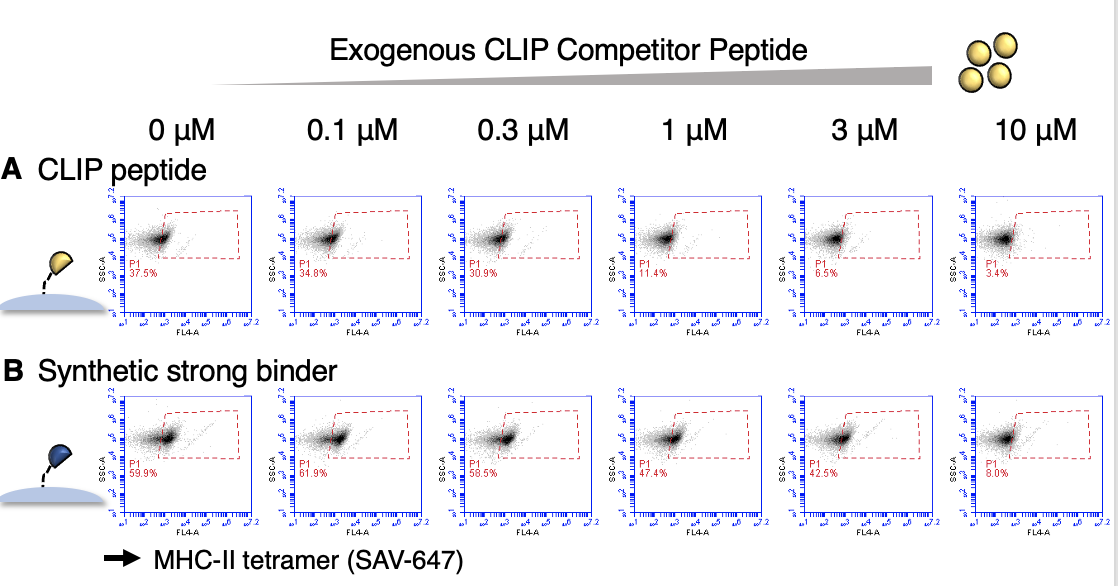

**Supplemental Figure 5. Competition with exogenous competitor peptide.** Binding of tetramerized HLA-DP401 to **A)** CLIP_81-101_ peptide or **B)** synthetic strong binding peptide expressed on yeast, and with exogenous CLIP_81-101_ peptide added as a competitor, from 0 μM to 10 μM.
